# Niche maintenance of germline stem cells in *C. elegans* males

**DOI:** 10.1101/428235

**Authors:** Sarah L. Crittenden, ChangHwan Lee, Ipsita Mohanty, Sindhu Battula, Judith Kimble

## Abstract

Stem cell maintenance by niche signaling is a common theme across phylogeny. In the *Caenorhabditis elegans* gonad, the broad outlines of germline stem cell (GSC) regulation are the same for both sexes: GLP-1/Notch signaling from the mesenchymal Distal Tip Cell (DTC) niche maintains GSCs in the distal gonad of both sexes (Austin and Kimble 1987), and does so via two key stem cell regulators, SYGL-1 and LST-1 (Kershner *et al*. 2014). Most analyses of niche signaling and GSC regulation have focused on XX hermaphrodites, an essentially female sex making sperm in larvae and oocytes in adults. Here we focus on XO males, which are sexually dimorphic in all tissues, including the distal gonad. The architecture of the male niche and the cellular behavior of GSCs are sex-specific. Despite these differences, males maintain a GSC pool similar to the hermaphrodite with respect to size and cell number and the male GSC response to niche signaling is also remarkably similar.

## INTRODUCTION

Niches maintain stem cells, prevent differentiation and provide a protective environment. Niches also have an architecture that is characteristic for any given tissue. This architecture – the positions and cellular contacts between niche and stem cells – is likely critical for stem cell regulation (O'Brien and Bilder 2013; Degirmenci *et al*. 2018). The *Caenorhabditis elegans* niche for germline stem cells (GSCs), one of the first niches identified (Kimble and White 1981), is sexually dimorphic and differs in number and position of niche cells between the sexes. The shape of the germline and cell cycle properties of the germ cells are also sexually dimorphic (Morgan *et al*. 2010). By contrast, key GSC regulators are similar in the two sexes: these include GLP-1/Notch signaling from the niche (Austin and Kimble 1987) and LST-1 and SYGL-1, which are the key targets of GLP-1/Notch signaling and fully account for the effect of niche signaling (Kershner *et al*. 2014). Therefore, *C. elegans* is poised to investigate the fundamental question of how cellular architecture of the niche region affects the size and shape of the niche response.

*C. elegans* GSCs reside in the “progenitor zone” region of the gonad in both sexes (Fig 1A–B). They are maintained there by a mesenchymal niche, the distal tip cell (DTC). Hermaphrodite gonads possess a progenitor zone at the end of each of the two gonadal arms, whereas male gonads have a single progenitor zone at the end of its single arm. Each arm, regardless of gender, contains hundreds of germ cells that are patterned along the distal-proximal axis, from GSCs at one end to differentiating gametes at the other. Hermaphrodites make sperm as larvae and oocytes in adults, whereas males make sperm continuously. Until overt gametogenesis, germ cells reside at the periphery of their gonadal arm and connect to a central rachis by cytoplasmic bridges (Hirsh *et al*. 1976; Amini *et al*. 2014; Seidel *et al*. 2018).

**Fig. 1.**
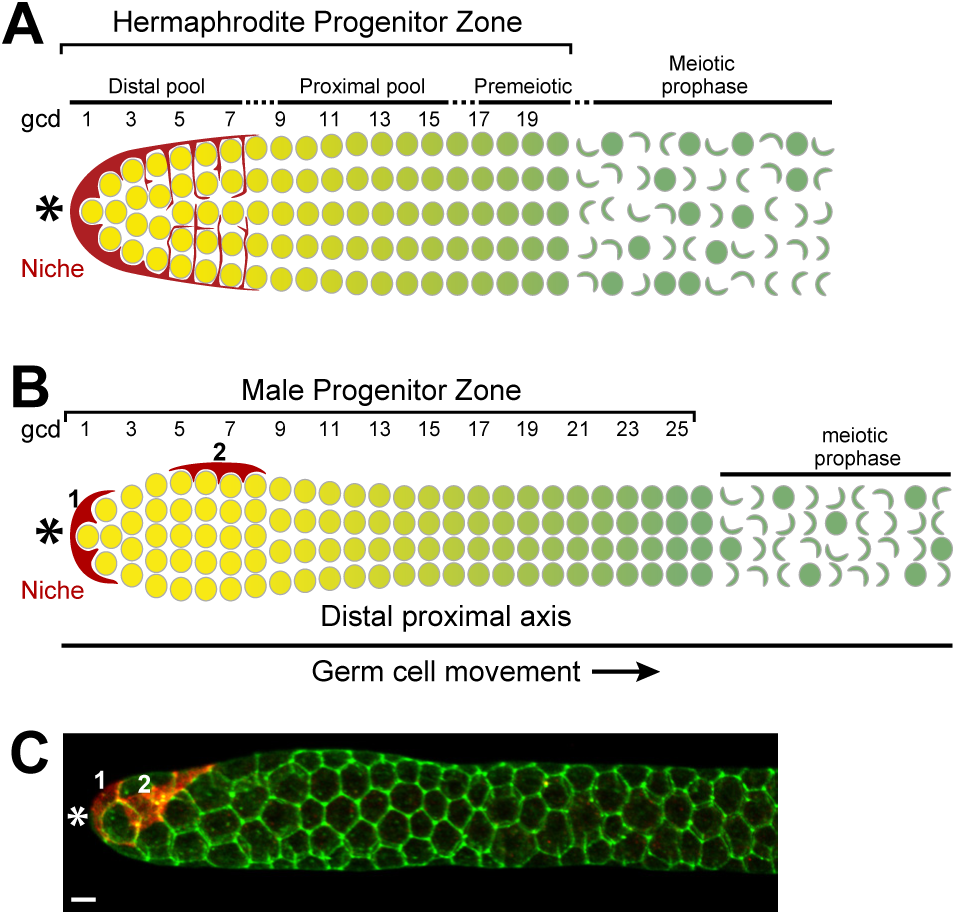
Introduction. (A) Diagram of hermaphrodite distal gonad. The progenitor zone contains a distal pool of naïve, undifferentiated cells (yellow) and a proximal pool of cells that are differentiating (graded green). As germ cells leave the proximal pool, they enter meiotic prophase (darker green). The DTC caps the distal rows of germ cells and has a plexus of processes that contact most cells in the distal pool. (B) Diagram of male distal gonad. The male progenitor zone extends further from the distal end, but contains the same number of germ cells as the hermaphrodite. Two DTCs are present in the male gonad. (C) Male distal gonad. DTCs are marked with *Parg-1*::myrGFP (red), germ cells are stained with anti-GLP-1 (green). DTCs are distal and lateral, GLP-1 is present in all cells in progenitor zone. Asterisks mark distal end of the germline, 1 and 2 mark male DTCs. Scale bar, 4 μm.

Both niches and progenitor zone are sexually dimorphic (Fig. 1A–B). The niche is composed of a single DTC in hermaphrodites, typically residing at the end of each progenitor zone, whereas it consists of two DTCs in males, with one positioned at the end of the progenitor zone and one on the dorsal side. The male progenitor zone often bulges right at the distal end, but overall is longer and thinner than in hermaphrodites (Fig. 1 A–B). Male germ cells cycle faster (Morgan *et al*. 2010) and progress through meiotic prophase more quickly (Jaramillo-Lambert *et al*. 2007). Germ cells normally fold into the center of the arm in adult hermaphrodites, but not in males (Raiders *et al*. 2018; Seidel *et al*. 2018). Finally, males are more sensitive to decreased GLP-1/Notch signaling than hermaphrodites.

Most detailed analyses of the DTC niche and its GSC response have focused on adult hermaphrodites. Here the DTC maintains a distal pool of germ cells in a stem cell-like state and a proximal pool of germ cells that progressively differentiate (Fig. 1A). The DTC and its processes extend around the distal-most germ cells, forming a plexus that correlates roughly with the distal pool (Cinquin *et al*. 2010; Byrd *et al*. 2014; Lee *et al*. 2016) and that contacts most of those germ cells (Lee *et al*. 2016). Cells in the distal pool express self-renewal markers, LST-1 and SYGL-1 (Kershner *et al*. 2014; Shin *et al*. 2017), but lack differentiation markers (Cinquin *et al*. 2010; Fox and Schedl 2015). By contrast, cells in the proximal pool lack self-renewal markers (Shin *et al*. 2017), but increasingly express differentiation markers, including both meiotic and sexual fate regulators (Hansen *et al*. 2004; Thompson *et al*. 2005; Cinquin *et al*. 2010; Fox and Schedl 2015; Brenner and Schedl 2016). Moreover, cells in the distal pool are essentially uniform in terms of cell cycle and respond similarly to cell cycle arrest (Crittenden *et al*. 2006; Cinquin *et al*. 2010; Rosu and Cohen-Fix 2017), whereas the more proximal cells undergo one or two mitotic divisions as they progressively differentiate and enter meiosis (Crittenden *et al*. 2006; Maciejowski *et al*. 2006; Cinquin *et al*. 2010; Fox and Schedl 2015; Rosu and Cohen-Fix 2017) (Fig. 1A). While we know male GSCs are maintained by DTCs in a progenitor zone, many features have not yet been characterized, including the extent of DTC processes, distal and proximal germ cell states and the molecular response to GLP-1/Notch signaling.

The molecular response to GLP-1/Notch signaling has been assessed in hermaphrodites (Lee *et al*. 2016; Shin *et al*. 2017). Whereas the receptor is found throughout the progenitor zone (Fig. 1C) (Crittenden *et al*. 1994; Crittenden *et al*. 2006; Cinquin *et al*. 2010), its transcriptional response is restricted to the distal stem cell-like pool and is steeply graded, with the highest response in cells closest to the DTC (Lee *et al*. 2016). The *sygl-1* and *lst-1* mRNAs and proteins are similarly restricted to the distal GSC pool (Kershner *et al*. 2014; Lee *et al*. 2016; Shin *et al*. 2017). This restriction is crucial to the size of the pool (Shin *et al*. 2017). While it is clear that GLP-1/Notch signaling maintains male GSCs, its transcriptional response and control over SYGL-1 and LST-1 expression in males is not yet known.

In this paper, we investigate the male niche and progenitor zone to ask whether there is sex-specific architecture of DTCs and whether this affects patterning in the progenitor zone. Male DTCs not only differ in number and position, but we find that they also differ in extent of contact with germ cells. In spite of this, we find that males have a distal GSC pool that is similar to that of hermaphrodites in size and number of cells, a similar transcriptional response to GLP-1/Notch signaling and a similar pattern of LST-1 and SYGL-1 protein expression. The extents of molecular patterns are very similar along the distal-proximal axis of gonad. Thus, in spite of different architectures and characteristics of both signaling and receiving cells, GLP-1/Notch output is nearly identical in the two sexes.

## Materials and Methods

### Strains and worm maintenance

*C. elegans* were maintained by standard techniques and grown at 20°C unless otherwise noted. Strains used were: N2, DG627 *emb-30(tn377ts)*, PD4443 *ccIs4443[Parg-1::GFP + dpy-20(+)]IV* (Kostas and Fire 2002), JK5764 *qSi361[Parg-1::myrGFP::tbb-2] IV*, JK5929 *lst-1(q1004) I* (Shin *et al*. 2017), JK6002 *sygl-1(q1015) I* (Shin *et al*. 2017), EG8081 *unc-119(ed3) III; oxTi177 IV* (Frøkjær-Jensen *et al*. 2014). Some strains were provided by the CGC, which is funded by NIH Office of Research Infrastructure Programs (P40 OD010440).

### Male DTC transgenes and scoring

For male DTC architecture we used strains containing either *Parg-1*::GFP (PD4443, *ccIs4443 IV*, (Kostas and Fire 2002)) or *Parg-1*::myrGFP (JK5764; *qSi361(IV)). Parg-1::myr* GFP strain is a mos insertion of pJK2010 (Parg-1::GFP with a myristoylation sequence inserted at the N-terminus of GFP). pJK2010 was created using Gibson assembly of 6.1 kb of genomic DNA upstream of the *arg-1* start site, myristoylated GFP from plasmid pJK1709 and the mos insertion plasmid pCFJ151 (Frøkjær-Jensen *et al*. 2014). The overall GFP expression pattern is very similar in PD4443 and JK5764, although GFP is membrane associated in JK5764, as expected, and levels appear lower, consistent with this being a single copy insertion. Animals were scored both live and fixed with similar results. Live animals were staged, mounted on agarose pads with either levamisole, 0.1 *μ*m Polybeads (Polysciences) or both and imaged on either a Zeiss LSM510 or Leica SP8 confocal microscope.

### *emb-30* assay

The *emb-30* assay was done as described previously (Cinquin *et al*. 2010). DG627 *(emb-30(tn377ts))* animals were grown at 15 °C until 36 h past L4 then shifted to restrictive temperature by moving plates to a 25 °C incubator or by using an EchoTherm timed incubator (Torrey Pines Scientific; incubator temperature shift occurred in less than 3 min). Animals at 15 °C were examined at 36 h past stage L4. Gonads were dissected, fixed and stained for anti-PH3, anti-GLD-1 and DAPI after the specified temperature regime. The length of the progenitor zone was measured in cell diameters and microns. To estimate the number of cells in the distal pool, we manually counted the number of M-phase arrested cells distal to the progenitor zone/transition zone boundary using the multipoint tool in FIJI (Schindelin *et al*. 2012). We consider this number a rough estimate since most germlines contained some fragmented nuclei, presumably a result of extended arrest, and *emb-30* mutants at permissive temperature do not have a wild-type number of progenitor zone cells (Cinquin *et al*. 2010). We excluded samples in which the majority of nuclei were fragmented because they could not be reliably counted.

### Immunohistochemistry

Males and hermaphrodites containing V5 tagged LST-1 or SYGL-1 or *Parg-1*::myrGFP were picked as L4s and gonads were dissected 22–26 hours later (A24 adults). Gonads were extruded, fixed and stained using standard protocols (Crittenden *et al*. 2017). Animals were washed off plates in ~ 200 *μ*l PBS containing 0.1% Tween 20 (PBSTw) and 0.25mM levamisole and pipetted into a multiwell dish. They were then cut behind the pharynx to extrude the gonad and then pipetted into a 1.5 ml microfuge tube and 4% paraformaldehyde was added for 10 minutes at room temperature. After centrifugation at 1500 rpm for 30 seconds, excess liquid was removed and gonads were permeabilized with 100 ul PBSTw containing 0.5% BSA (PBSBTw) for 5 minutes at room temperature. Centrifugation was repeated and gonads were blocked in PBSBTw for 30 minutes at room temperature. Primary antibodies were added and tubes were placed at 4°C overnight. Mouse anti-V5 (Bio-Rad) was used at 1:1000-1:5000, Mouse anti-GFP (3E6, Invitrogen) was used at 1:200, Rabbit anti-GLP-1(LNG) (Crittenden *et al*. 1994) was used at 1:20. Secondary antibodies (Alexa 488, Alexa 555 and Alexa 647; Jackson ImmunoResearch) were used at 1:1000 in PBSBTw and incubated for 1 hour at room temperature on a rotating rack shielded from light, followed by 3 washes in PBSBTw. Finally, samples were mounted in 10 *μ*l ProLong Gold (ThermoFisher) and covered with a 22×22 coverslip, allowed to cure overnight to several days and imaged on a Leica SP8 confocal microscope.

### smFISH and analysis

We used the previously described smFISH protocol, imaging parameters and analysis tools with minor modifications (Lee *et al*. 2016; Lee *et al*. 2017). Males and hermaphrodites, 24 hours after mid-L4, were dissected in ~ 200 *μ*l PBS, 0.1% Tween 20 (PBSTw) containing 0.1% tricaine and 0.01% tetramisole. Dissected worms were pelleted at 1500 rpm for 30 seconds, fixed in 1 ml PBSTw, 3.7% formaldehyde, pelleted, washed in PBSTw and permeabilized in PBS containing 0.1%Triton X100 for 10 minutes at room temperature. Samples were washed 2 times and resuspended in 1 ml 70% ethanol and stored at 4°C overnight or up to one week. Samples were washed, hybridized with probes *(sygl-1* intron, 48 unique oligonucleotides, QUASAR 570; *sygl-1* exon, 31 unique oligonucleotides, CALFLUOR 610; and/or *let-858* intron, 48 unique oligonucleotides, CALFLUOR 610; Biosearch Technologies) overnight at 37°C, protected from light. Following hybridization, samples were washed at room temperature, incubated with 1 ml wash buffer containing 1 *μ*g/ml DAPI for 30 minutes, washed 2 additional times then mounted in ProLong Gold and allowed to cure overnight to several days before imaging.

Images were acquired using a Leica SP8 confocal microscope and quantitated using MATLAB essentially as described (Lee *et al*. 2016).

### LST-1 and SYGL-1 scoring

LST-1:V5 and SYGL-1:V5 (Shin *et al*. 2017) were quantitated in confocal stacks using FIJI/ImageJ. Briefly, a line (linewidth = 50 pixels) was drawn along the distal proximal axis of the germline. Pixel intensity was measured using the plot profile function. Intensities were copied into MATLAB, averaged and graphed. An N2 control was used to quantitate background for anti-V5.

## RESULTS

### Architecture of the DTC niche in males

The single male gonad contains two DTCs (Kimble and Hirsh 1979). One DTC typically caps the distal end, while the other is typically dorsal (Figs. 1B, 2A–B) (Kimble and White 1981). Both DTCs are required to maintain the normal number of cells in the progenitor zone and either DTC can partially maintain the progenitor zone (Kimble and White 1981; Morgan *et al*. 2010). Architecture of the hermaphrodite DTC (hDTC) has been well described (Byrd *et al*. 2014; Linden *et al*. 2017). The DTC cap has extensive membrane contact with the distal-most 3–4 rows of germ cells (Byrd *et al*. 2014; Lee *et al*. 2016; Linden *et al*. 2017). Adult hDTCs extend processes along the germline surface and also intercalate between germ cells; the extent of this plexus of extensive germ cell contacts correlates well with the distal GSC pool (Byrd *et al*. 2014; Lee *et al*. 2016). Long surface processes can also extend beyond the progenitor zone.

**Fig. 2.**
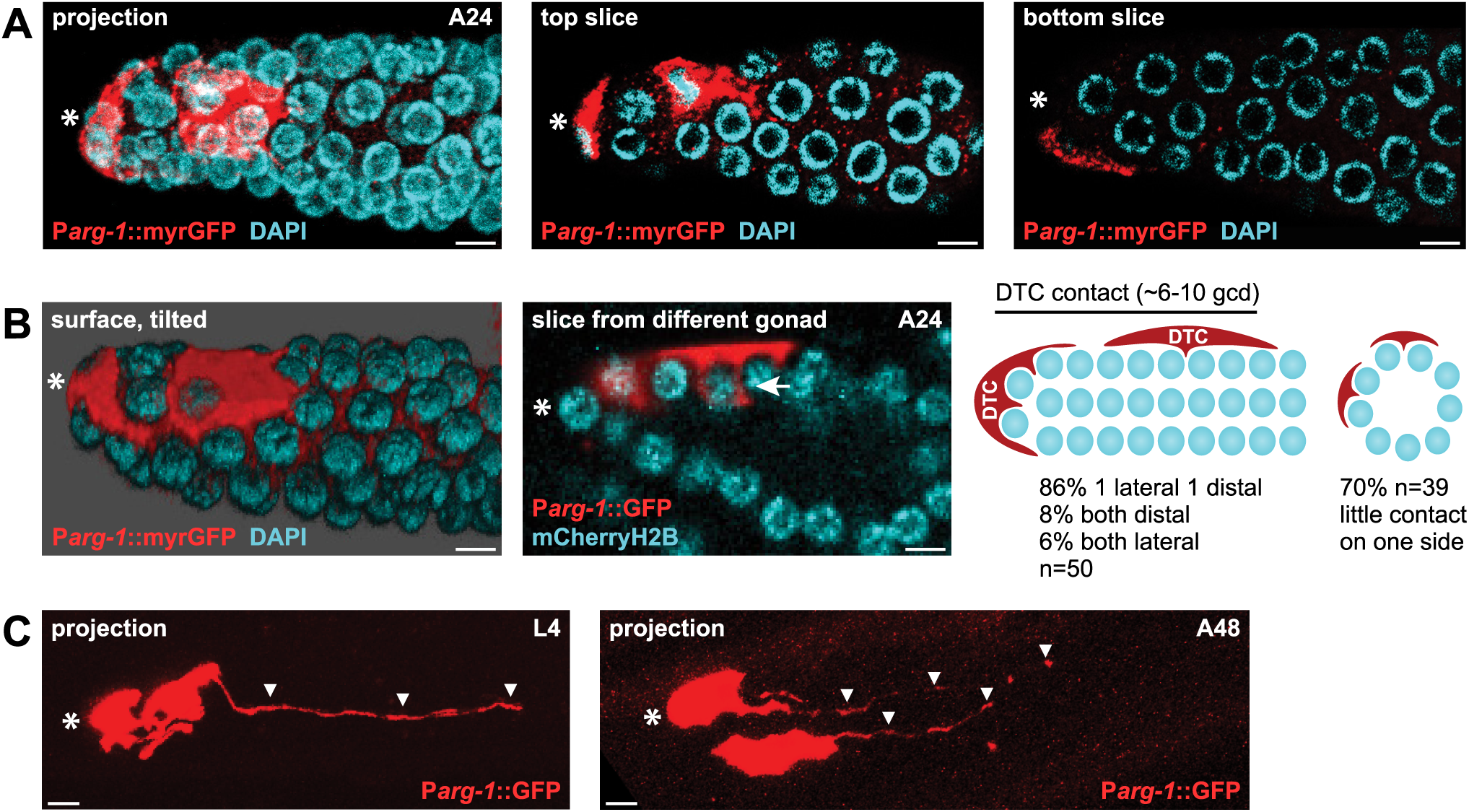
Architecture of male DTCs. DTCs are red, nuclei are cyan. Distal end is marked with a white *. (A) Projection and slices (labeled in panels) from a confocal z stack of a representative adult (A24) male gonad. DTCs are distal and lateral and can have processes. In the bottom slice a number of germ cells lack detectable DTC processes. (B) Left: Surface rendering of gonad in (A), tilted to show another view of DTC contact with germ cells. Middle: a different male gonad showing short intercalating processes around germ cells adjacent to a lateral DTC cap. About 85% of gonads (n=50) have short processes that embrace adjacent germ cells (white arrow). Right: Diagram of DTC position and percent gonads with various configurations and diagrammatic view of a cross section showing lack of detectable DTC contact on one side of the gonad. (C) L4 and A48 male gonads. DTC complexity does not seem to be a function of age in male gonads as it is in hermaphrodites (Byrd *et al*. 2014). We could see either more extensive processes or the more typical less extensive contact in both young and old gonads. Scale bars, 4 μm.

To examine the contact of male DTCs (mDTCs) with germ cells, we used animals carrying transgenes with the *arg-1* promoter driving either GFP (Kostas and Fire 2002; Zhao *et al*. 2007) or GFP containing a membrane-targeting myristoylation sequence (myrGFP) in mDTCs. We first scored the position of each mDTC along the distal proximal axis of the germline (Fig. 2A–B) in animals 24 hours past L4. Most gonads have one mDTC at the distal end (47/50) and one placed laterally from the end (43/50). In some cases, they reside next to each other, either at the end (4/50 or laterally (3/50). In live animals, the lateral mDTC is typically dorsal (90%, n=20). We next scored the extent of each DTC. As with the hDTC, the cell body of each mDTC forms a cap that has extensive contacts with 3–5 rows of germ cells (n=35; Fig. 2A–B); the two DTCs together extend 6–10 germ cell diameters (gcd) on average from the distal end (Fig. 2B). Similar to hDTCs, mDTCs cover the germ cell surface and extend short processes around them (Fig. 2A–B; 84%, n=50). In a few cases (16%, n=50), it was difficult to see short processes. Yet, in contrast to hDTCs, the mDTCs did not have an extensive plexus of processes; only 1/50 generated a plexus. In only some gonads (19/50), we found long superficial processes. Overall, we found that while some components of DTC architecture were similar in adult males and hermaphrodites, others differed. mDTC cap morphology was similar to hDTC cap morphology. Both have extensive contact with adjacent germ cells. However, adult mDTCs typically lack the extensive plexus and long external processes seen in adult hDTCs.

One striking difference between male and hermaphrodite niche architecture is that mDTC contacts are not uniform around the germline; most had little or no detectable contact on one side (~27/39 (~70%)) (Fig. 2A–B). Another difference is developmental: hDTC morphologies are distinct in L4s and young adults, developing the plexus soon after molting (Byrd *et al*. 2014; Linden *et al*. 2017). By contrast, mDTCs look similar in L4s and young adults, with long surface processes at the same frequency. However, mDTC processes tend to lengthen and gain processes after two days (n=3) (Fig. 2C).

In conclusion, the extent and morphology of processes is variable between sexes and between developmental stages. The male-specific architecture of the niche raises the question of whether there is also a male-specific pattern of GLP-1/Notch response in the germline.

### Males have a GSC pool adjacent to the niche

To ask how male germ cells are patterned within the progenitor zone, we used the *emb-30* assay (Fig. 3) (Cinquin *et al*. 2010). In hermaphrodites, this assay defined the distal GSC pool. The assay used a temperature-sensitive mutant of *emb-30*, which is required for the metaphase to anaphase progression (Furuta *et al*. 2000). Without *emb-30* activity, germ cell cycling stops and distal to proximal movement halts, allowing germ cells to reveal their developmental state in place (Fig. 3A). In hermaphrodites, a distal pool remained arrested and undifferentiated, expressing little of the differentiation marker GLD-1, while the proximal pool of germ cells progressively differentiated, forming crescent shaped DNA typical of early meiotic prophase and expressing high GLD-1.

**Fig. 3.**
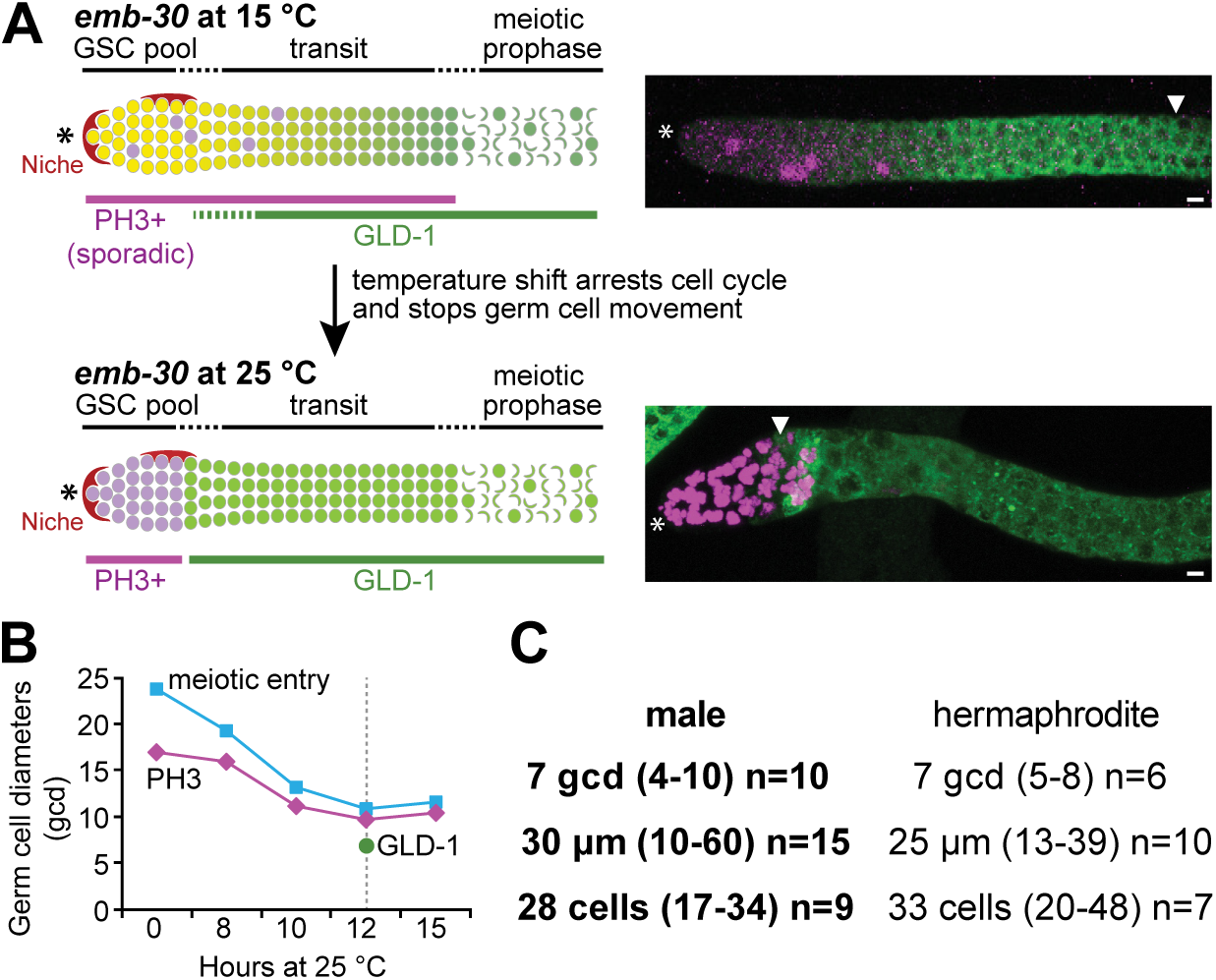
GSC pool. (A) *emb-30* assay. Distal end of gonad is marked with *, progenitor zone boundary is marked with white triangle. Left: Diagram of experimental design. Right: Representative confocal projections of distal germlines from *emb-30* males shifted to restrictive temperature for the indicated times. Germlines are stained with anti-GLD-1 (green) and anti-PH3 (magenta). (B) As germ cells arrest, the number of PH3-positive nuclei increases with time and the position of both PH3-positive (magenta line) and meiotic entry (cyan) becomes more distal with time, plateauing at about 12 hours. Entry into meiosis occurred at about 11 gcd (5–17, n=28); the region distal to this contained ~45 germ cells on average (26–82, n=19), similar to hermaphrodites (Cinquin *et al*. 2010). The GLD-1 boundary (green dot) after 12 hours at restrictive temperature is indicated. (C) GSC pool characteristics were measured at the 12 hour time point. Males and hermaphrodites were done in parallel.

**Fig. 4.**
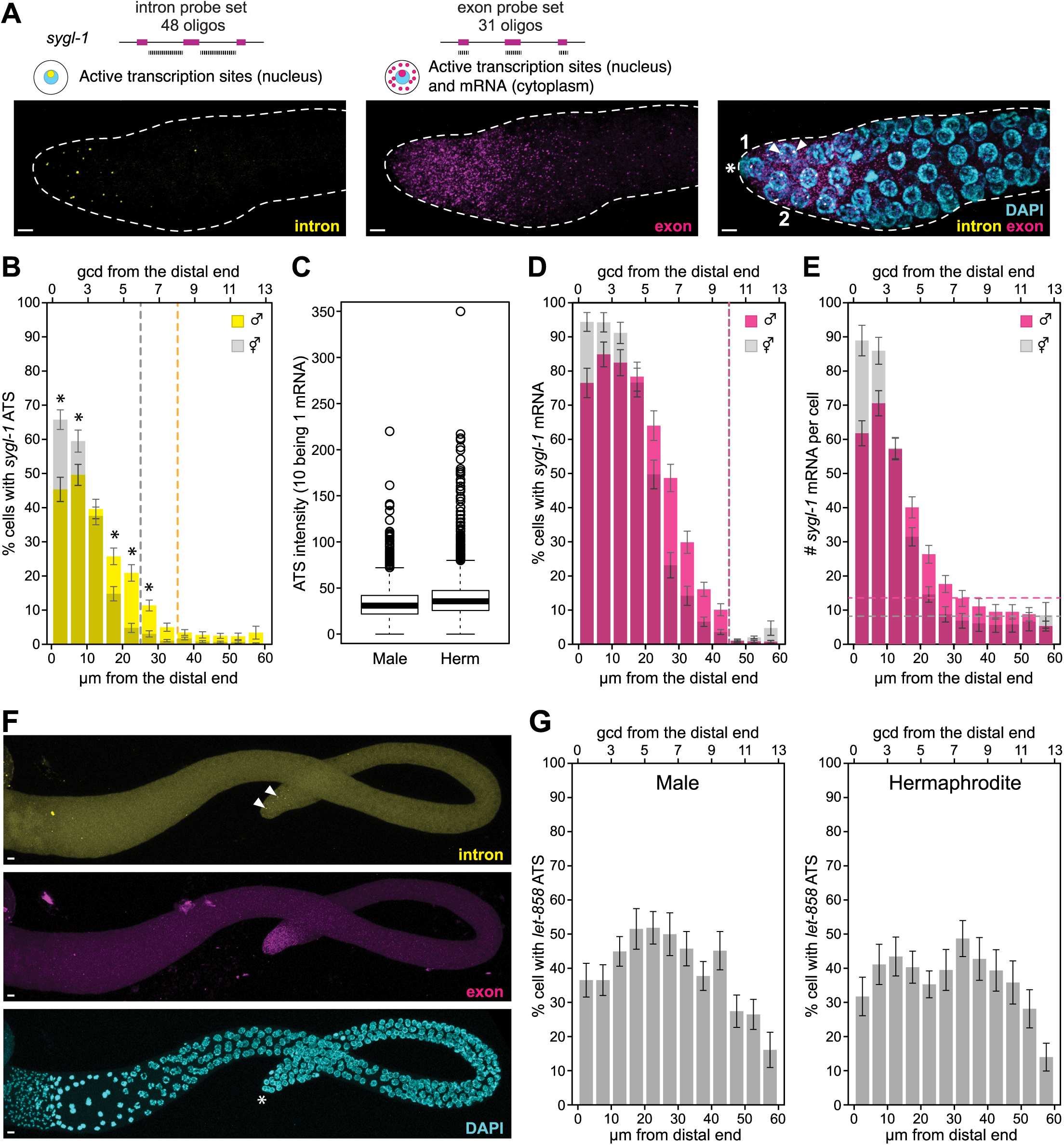
*sygl-1* smFISH. (A) Representative images of distal end of a male gonad showing intron probe, exon probe and merge of intron and exon probes with DAPI. smFISH probes and expected subcellular distribution for exon and intron probes are diagrammed above images. Distal end is marked with asterisk and positions of DTC nuclei are marked 1 and 2 in the merged panel. Two ATS are marked with white arrowheads in the merged pane. (B-E) MATLAB quantitation of male and hermaphrodite gonads done in parallel. Overall pattern is similar in males and hermaphrodites. Males n=64, hermaphrodites n=53. Error bars: Standard error of the mean. (B) % cells with ATS above baseline. Male (yellow bars) and hermaphrodite (grey bars). Asterisks indicate significant differences. p-value for paired t-test is < 0.01. Males have a lower % cells with ATS in the distal-most rows of germ cells. Dotted lines mark the boundaries in males (yellow) and hermaphrodites (grey) where percent cells with ATS drops below baseline levels. (C) Signal intensity of ATS in male (left) and hermaphrodite (right). Mean for males is 34.4, for hermaphrodites is 41.1. p-value for t-test is 6.4x10^-15^. The signal intensity of 1 mRNA is 10, so males on average have ~0.5 mRNA less per ATS than hermaphrodites. (D) Percent cells with mRNA above baseline. Male (pink bars) and hermaphrodite (grey bars). Dotted lines mark the boundaries in males (pink) and hermaphrodites (grey) where % cells with mRNA drops below baseline levels. Males and hermaphrodites have the boundary at the same point. (E) Number of mRNA above baseline per cell. Male (pink bars) and hermaphrodite (grey bars). Dotted lines mark average baseline levels for male (pink) and hermaphrodites (grey). (F) *sygl-1* smFISH on entire male germline. Hermaphrodites have strong, GLP-1/Notch independent transcription in the proximal germline (Kershner *et al*. 2014; Lee *et al*. 2016). Males do not. (G) MATLAB quantitation of ATS detected using *let-858* exon probes. Males n=24, hermaphrodites n=20. We do not detect a sex-specific difference in percentage of cells with ATS using this control probe, in contrast to what was seen with the *sygl-1* probe (Fig. 4). We also do not see graded *let-858* ATS in the GSC pool of either sex.

To reveal the male distal GSC pool and estimate its size, we used a time course to learn when distal and proximal pools resolved (Fig. 3B). We raised *emb-30(ts)* males at permissive temperature until the L4 stage, then switched them to restrictive temperature and scored the position of early meiotic prophase and PH3-positive cells after 0, 8, 10, 12 and 15 hours. After 12 hours, the boundary stabilized between undifferentiated and differentiated cells (Fig. 3B). We then stained for GLD-1 after 12 hours at restrictive temperature and measured the boundary between low and high GLD-1 levels; this analysis identified a distal pool of arrested germ cells with low GLD-1 and a proximal pool with high GLD-1. The extent of the distal pool extended between 4 and 10 (average 7, n=10) germ cell diameters (gcd) from the distal end and contained between 17 and 34 (average 28, n=9) germ cells. This extent and cell number were similar to those in hermaphrodites done in parallel (Fig. 3C). The actual number of germ cells in the GSC pool is likely higher than the estimate from *emb-30* experiments (Cinquin *et al*. 2010); in wild-type male germlines there are roughly 35–50 germ cells in the distal-most 6–8 rows (Morgan *et al*. 2010; also Fig. 5B). We conclude that the male progenitor zone, like that of hermaphrodites, possesses a distal pool of germ cells maintained in a naïve state as well as a proximal pool of cells primed to differentiate. Moreover, our estimate of both the extent and number of cells in the distal pool is similar in the two sexes.

**Fig. 5.**
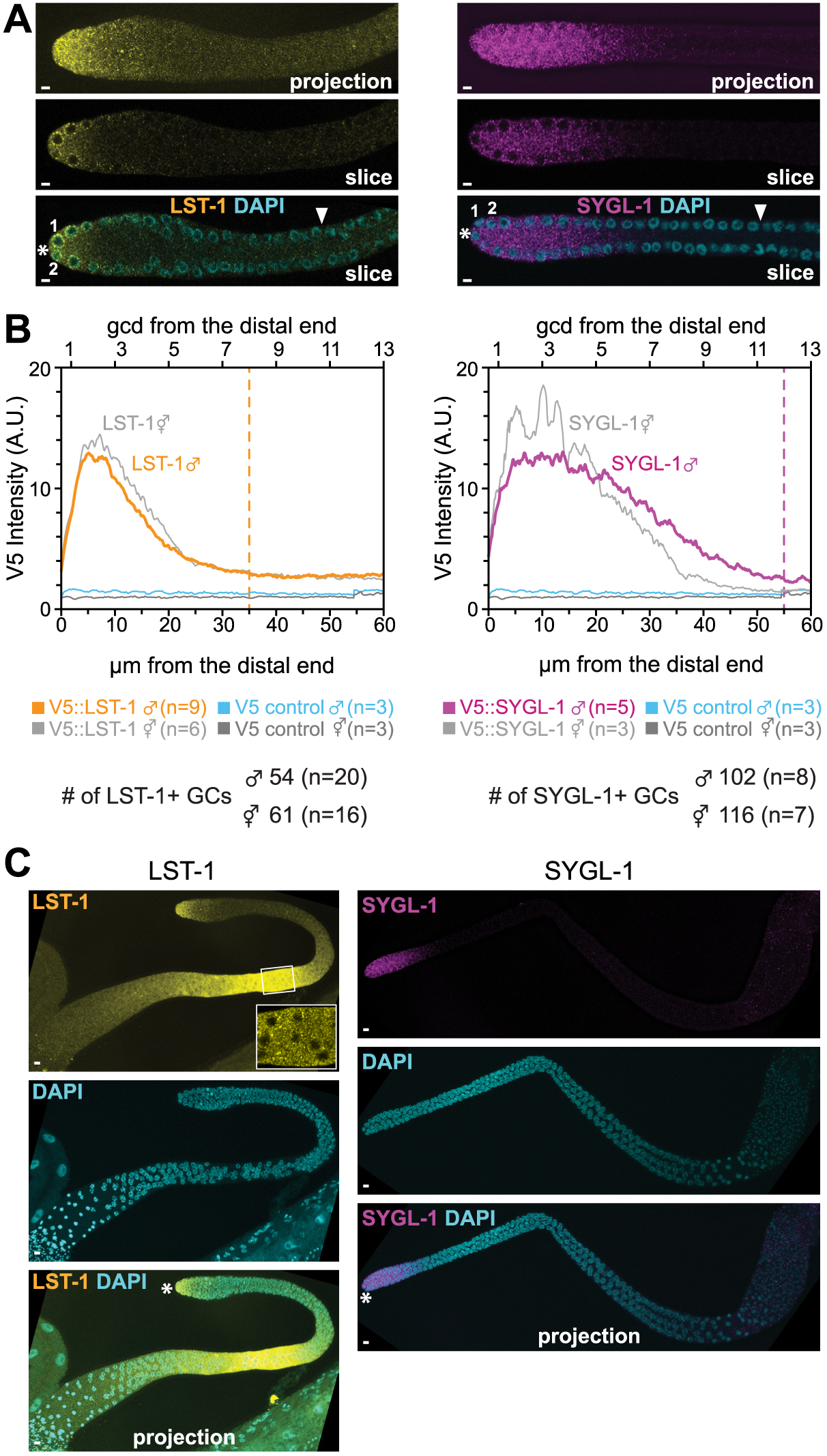
LST-1 and SYGL-1 proteins are restricted to the male GSC pool in the distal germline. (A) LST-1 (left) and SYGL-1 (right) staining in distal male gonads. Top panels: single channel maximum intensity confocal projections. Middle panels: single channel, single confocal slice through middle of gonad. Bottom panels: same as middle panel merged with DAPI. Distal end is marked with asterisk, 1 and 2 indicate positions of DTC nuclei. (B) ImageJ quantitation of summed projections of LST-1 and SYGL-1 stained gonads. LST-1 (left), SYGL-1 (right). Controls (blue: male, grey: hermaphrodite) were N2 animals stained with anti-V5 in parallel with V5-tagged LST-1 and SYGL-1 lines. Dotted lines indicate the average position where either LST-1 or SYGL-1 dropped to background levels when scored by eye. Number of germ cells that are LST-1 or SYGL-1 positive in male or hermaphrodite germlines are shown below the graphs. (C) Projected confocal z stacks of entire adult male germlines stained for either LST-1 (left) or SYGL-1 (right). Males have proximal LST-1 (left) but not SYGL-1 (right). LST-1 is predominantly cytoplasmic in the proximal region (inset, single slice from proximal region). Hermaphrodites have proximal SYGL-1 but not LST-1 (Shin *et al*. 2017).

### Male GLP-1/Notch transcriptional response is graded and restricted to the distal GSC pool

We next analyzed the transcriptional response to GLP-1/Notch signaling in males to ask if it had male-specific characteristics. In hermaphrodites, this response is steeply graded and restricted to the GSC pool region (Lee *et al*. 2016). The gradient was unexpected since essentially all germ cells in the pool contact the DTC (Lee *et al*. 2016), all express receptor and all appear to be in a similar naïve state (Cinquin *et al*. 2010; Fox and Schedl 2015). Interestingly, hermaphrodite L4s had a similar response despite the difference in niche morphology (Byrd *et al*. 2014; Lee *et al*. 2016).

To test the male response, we used *sygl-1* smFISH to identify active transcription sites (ATS) and cytoplasmic mRNAs, using a combination of intron-specific and exon-specific probes (Fig. 4A), as done previously. We identified ATS based on established criteria (Lee *et al*. 2016; Lee *et al*. 2017), including colocalization of exon and intron probes as well as nuclear location, assessed by DAPI. We identified mature, cytoplasmic mRNAs based on signal from exon probes alone (Lee *et al*. 2016) (Fig. 4A). We immediately noticed that both ATS and mRNAs were restricted to the distal end of the progenitor zone, as in hermaphrodites; the extent correlates roughly with the GSC pool. We also noticed that, in males, both ATS and mRNA appeared dimmer at the distal end. In many germlines we saw ATS and mRNA away from the lateral DTC. Hermaphrodites transcribe *sygl-1* in a GLP-1-independent manner in the proximal germline, likely as a maternal RNA (Shin *et al*. 2017); this proximal expression was not seen in males (Fig. 4F).

To quantitate the *sygl-1* smFISH results, we used a modified MATLAB code developed previously to analyze smFISH in hermaphrodites (Fig. 4B) (Lee *et al*. 2016; Lee *et al*. 2017). The MATLAB results confirmed what we initially noticed and gave us a quantitative view of the transcription pattern. Overall, the pattern is similar in male and hermaphrodite germlines: both ATS and mRNAs are restricted to the distal end and ATS probability is graded with distal-most cells more likely to have an ATS (Fig. 4B). ATS intensity is similar between the sexes (Fig. 4C). The number of *sygl-1* mRNAs per cell is also graded and, as in hermaphrodites, its pattern is expanded proximally (~10 gcd) compared to the ATS pattern (~8 gcd) (Fig. 4D). There are two differences between the sexes. First, the percentage of cells with ATS was lower at the distal-most end (Fig. 4B) as were the percentage of cells with *sygl-1* mRNA and the number of *sygl-1* mRNAs per cell (Fig. 4D–E). The second difference is that the ATS region extends further from the distal end and (Fig. 4B). This expansion is small, about 1 cell diameter, but statistically significant and visible by eye. As a control, we assayed *let-858*, which is ubiquitously expressed (WormBase), and found its distribution and levels identical in males and hermaphrodites in the distal region of the gonad (Fig. 4G).

We conclude that the pattern of the GLP-1/Notch transcriptional response is similar in the two sexes and that surprisingly, the sex-specific niche architecture in males does not have a major effect on this pattern.

### LST-1 and SYGL-1 proteins are restricted to the GSC pool in the male distal germline

Last, we analyzed the expression of LST-1 and SYGL-1 proteins. In hermaphrodites, both are expressed in the GSC pool and their extent correlates well with extent of the GSC pool. To ask if LST-1 and SYGL-1 are similarly restricted in males, we stained epitope-tagged LST-1 and SYGL-1 (Shin *et al*. 2017) in adult male germlines. LST-1 and SYGL-1 are found in the GSC pool region in male germlines: LST-1 is abundant in the distal-most ~5–6 gcd and is detectable in the distal-most ~8 gcd, SYGL-1 is abundant in the distal-most ~7–8 gcd and is detectable in the distal-most ~12 gcd (Fig. 5 A–B). In the distal germ cells, LST-1 is both cytoplasmic and associated with perinuclear granules, while SYGL-1 is cytoplasmic in both sexes (Fig. 5A). Remarkably, their patterns and levels of expression were similar in the distal germlines of males and hermaphrodites (Fig. 5B). This is particularly striking because *sygl-1* transcription in males is lower than in hermaphrodites in the distal 4 rows of the male germline, yet SYGL-1 protein abundance is comparable in the two sexes. In addition to expression in distal germ cells, LST-1 is found in proximal male germ cells in the pachytene region of the gonad (Fig. 5C). In this region, LST-1 is predominantly cytoplasmic (Fig. 5C, inset). SYGL-1 is not present proximally (Fig. 5C), consistent with the lack of *sygl-1* RNA in this region (Fig. 4F).

## DISCUSSION

In this paper, we compare key features of the hermaphrodite and male niche regions to ask if niche architecture and sex-specific germ cell characteristics modulate GSC response to niche signaling. In addition to male-specific DTC number and positioning, we find that adult male DTCs do not form extensive processes as they do in adult hermaphrodites. In spite of these differences in niche architecture, we find only minor sex-specific differences in the patterning of the progenitor zone and the molecular response to niche signaling.

The *C. elegans* male GSC pool shares many characteristics with the hermaphrodite GSC pool (Fig. 6), including number of cells in the pool (~30–50), extent along the distal proximal axis of the germline, extent of transcriptional response to GLP-1/Notch signaling, and level and extent of key GSC regulators, LST-1 and SYGL-1. Thus, GSC pool characteristics are shared in spite of sexually dimorphic germline and niche architecture, differences in cell cycle speed and differences in proteins that regulate differentiation.

**Fig. 6.**
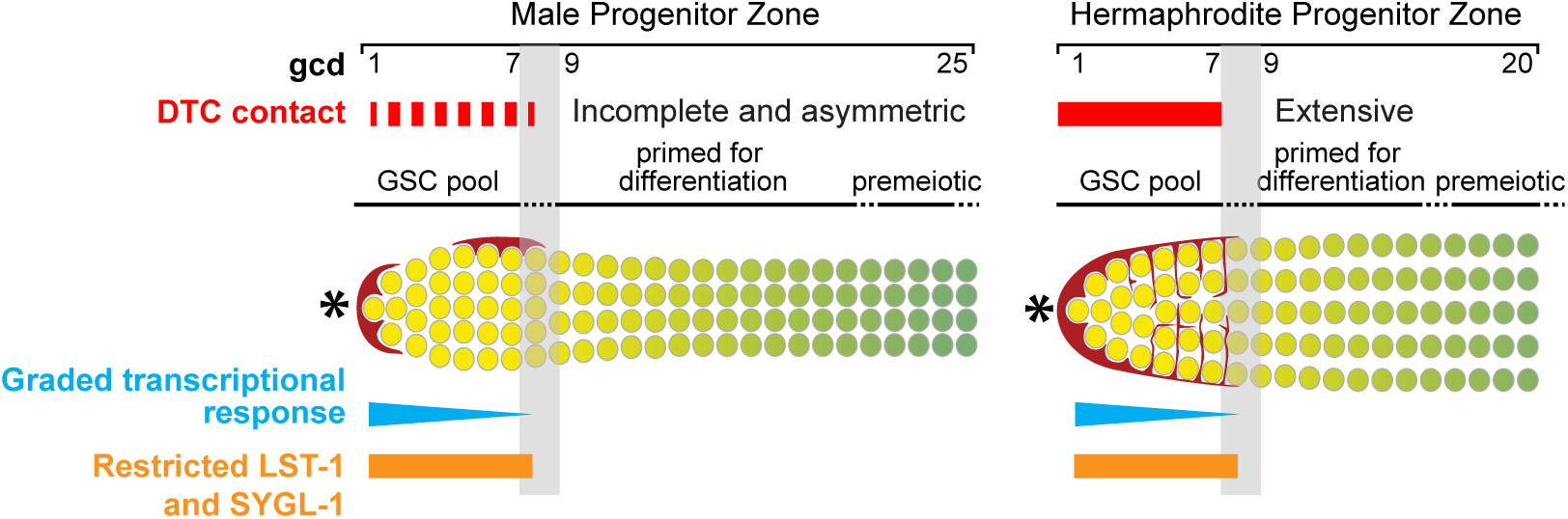
Summary of male GSC niche region and comparison to hermaphrodite. The male and hermaphrodite progenitor zones have very similar patterns in spite of different DTC architecture. Male DTCs have caps (red), hermaphrodite DTCs have a cap and extensive plexus surrounding the GSC pool (red, red lines between GSCs in diagram). Male DTC contact with germ cells is incomplete and asymmetric (dashed red bar) compared to the hermaphrodite (red bar). The response to GLP-1/Notch signaling is similar in males and hermaphrodites. The transcriptional response is restricted to and graded through the GSC pool (blue bars) and the key target proteins, LST-1 and SYGL-1, are restricted to the distal region of the germline with similar levels and extents in both sexes (orange bars).

While the male GSC pool shares many characteristics with the hermaphrodite GSC pool, there are some minor differences. The transcriptional response, both ATS and mRNA, is slightly lower in the distal-most rows of germ cells and is slightly increased in more proximal rows. Perhaps this indicates altered signaling dynamics due to male specific DTC architecture, ligands or downstream regulatory events. In contrast, SYGL-1 protein levels are similar in GSCs in both sexes. In addition, SYGL-1 protein is expanded slightly along the distal-proximal axis. Perhaps the additional DTC in males and the sex-specific characteristics of male germ cells and their regulators contribute to the slight expansion in males. Alternatively, the pattern may appear shorter in hermaphrodites due to folding in the oogenic hermaphrodite germline (Raiders *et al*. 2018; Seidel *et al*. 2018).

It is notable that very similar transcriptional responses are seen in gonads with very different DTC architectures. One component that is consistent in gonads from different stages and sexes is the distal cap region. Male gonads contain one distal and one dorsal DTC, each of which caps adjacent germ cells and contains minimal processes. During hermaphrodite larval development, the DTC caps adjacent germ cells, but does not have detectable processes (Byrd *et al*. 2014; Linden *et al*. 2017; Pekar *et al*. 2017). The adult hermaphrodite DTC caps adjacent germ cells and also has an extensive plexus that contacts virtually all cells in the GSC pool (Byrd *et al*. 2014; Lee *et al*. 2016; Linden *et al*. 2017). The extensive contact between the cap and germ cells may facilitate increased signaling capability (Shaya *et al*. 2017), whereas the plexus and long processes have less extensive and likely labile contact with germ cells (Wong *et al*. 2013) and may not signal as effectively. There is evidence that DTC components other than the distal cap do have a function in the progenitor zone. In adult hermaphrodites, the processes appear to have a role in maintaining the progenitor zone (Linden *et al*. 2017) and in the males, both DTCs are required for normal progenitor zone size and each is sufficient to maintain the progenitor zone (Kimble and White 1981; Morgan *et al*. 2010). Perhaps the processes have roles other than activation of GLP-1/Notch signaling in maintaining the GSC niche. These could include positioning the GSCs, protecting them from contact with other tissues or providing signals important for robust proliferation.

The variation in niche architecture raises the question of how the molecular response to GLP-1/Notch signaling is shaped. The transcriptional response in a subset of the GSC pool occurs both with (adult hermaphrodite) and without (L4 hermaphrodites and males) detectable DTC contact. One simple model is that ligand produced by the DTCs is broadly distributed and in contact with most cells in the GSC pool, whether or not there are detectable processes. Ligand could be presented on processes such as cytonemes (Huang and Kornberg 2015), that are thin and difficult to image. Alternatively, ligand could be diffusible and able to contact all of the GSC pool. There are several ligands in *C. elegans* that lack a transmembrane domain (Chen and Greenwald 2004; Komatsu *et al*. 2008), including *dsl-1* which plays a role in Notch-mediated vulval patterning (Chen and Greenwald 2004). In addition, the transcriptional response is also graded. The gradient could be generated by graded ligand presentation or perhaps more proximal germ cells become refractory to GLP-1/Notch signaling as they move away from the distal end of the gonad, either because they lack a key component of the GLP-1/Notch transcriptional complex or have some as yet unidentified regulator that modulates the response. Visualization of nuclear GLP-1 NICD, other components of the signaling complex as well as GLP-1/Notch ligands should help clarify these models.

## Acknowledgements

We thank Brandon Taylor for cloning *Parg-1::myr* GFP and DTC discussions, Karla Knobel and Dana Byrd for initial observation of *Parg-1*::GFP in male DTCs, Laura Vanderploeg and Kim Haupt for help with figures, Tina Lynch and Kim Haupt for comments on the manuscript and Anne Helsley-Marchbanks for assistance with manuscript preparation. Funding was provided by the National Institutes of Health (GM069454 to JK) as well as the Stem Cell & Regenerative Medicine Center (SCRMC) within the UW School of Medicine and Public Health. JK is an investigator of the Howard Hughes Medical Institute.

